# Reduction of genotyping marker density for genomic selection is not an affordable approach to long-term breeding in cross-pollinated crops

**DOI:** 10.1101/2021.03.05.434084

**Authors:** Júlio César DoVale, Humberto Fanelli Carvalho, Felipe Sabadin, Roberto Fritsche-Neto

**Author notes:** Corresponding author Department of Crop Science, Federal University of Ceará, Fortaleza, Ceará, Brazil, Júlio César DoVale.

## Abstract

The selection of informative markers has been studied massively as an alternative to reduce genotyping costs for the genomic selection (GS) application. Low-density marker panels are attractive for GS because they decrease computational time-consuming and multicollinearity beyond more individuals can be genotyped with the same cost. Nevertheless, these inferences are usually made empirically using “static” training sets and populations, which are adequate only to predict a breeding program’s initial cycles but might not for long-term cycles. Moreover, to the best of our knowledge, none of these inferences considered the inclusion of dominance into the GS models, which is particularly important to predict cross-pollinated crops. Therefore, that reveals an important and unexplored topic for allogamous long-term breeding. To achieve this goal, we employed two approaches: the former used empirical maize datasets, and the latter simulations of long-term breeding cycles of phenotypic and genomic recurrent selection (intrapopulation and reciprocal). Then, we observed the reducing marker density effect on populations (mean, the best genotypes performance, accuracy, additive variance) over cycles and models (additive, additive-dominance, specific combining ability (SCA)). Our results indicate that the markers reduction based on different linkage disequili brium (LD) levels is viable only within a cycle and brings a significant decrease in predictive ability over generations. Furthermore, in the long-term, regardless of the selection scheme adopted, the more makers, the better because they buffer LD losses caused by recombination over breeding cycles. Finally, regarding the accuracy, the additive-dominant models tend to outperform the additive ones and perform similar to the SCA.

## INTRODUCTION

Genomic selection (GS), using high-density single-nucleotide polymorphisms (SNPs) chips, has been widely adopted to improve prediction ability and response to selection (Hayes *et al*., 2009; Crossa *et al*., 2010). However, these chips’ genotyping cost is currently prohibitive, especially for low profitability species (Hou *et al*., 2020), and programs in poor regions, such as those in Latin America, Africa, and Southwest Asia.

In this context, density reduction of markers has been evaluated massively as an alternative to reduce genotyping costs for GS applications. This purpose can be achieved based on different criteria, for instance, marker effects of effect (Sousa *et al*., 2019), positions (Zhang *et al*., 2015), preselection based on haplotype block analysis (Ma *et al*., 2016); genome-wide association studies (GWAS) approach in which significant markers are selected to fit a prediction model (Subedi *et al*., 2013), and by linkage disequilibrium (LD), where highly correlated markers are eliminated by redundancy. All strategies have advantages and disadvantages. Among them, the LD criteria seem to be more suitable because it reduces multicollinearity, which is an obstacle for highly dense SNP panel, not delivers noise in the marker effects (Xu, 2013), and the marker reduction process is independent of the phenotype, reducing the probability of overfitting.

Low-density marker panels are also attractive for breeding purposes because they decrease computational time-consuming, and then more individuals can be genotyped with the same cost (Gorjanc *et al*., 2015). With larger datasets of phenotyped and genotyped individuals, GS’s predictive ability may be increasingly driven by LD rather than by linkage information (Hickey *et al*., 2014). Furthermore, several studies claim that it is possible to substantially reduce the number of markers and maintain high predictive ability (Tayeh *et al*., 2015; Ma *et al*., 2016; Sousa *et al*., 2019; Al-Tobasei *et al*., 2020). Thus, the use of genotyping techniques with highly discriminative markers could be a viable alternative to current low-cost SNP array strategies, with the potential to increase the fraction of the genome captured in a cost-efficient manner (Elshire *et al*., 2011).

Nevertheless, in all these studies, it was possible to obtain an informative marker subset either with empirical or simulated data. The results are usually good because they consider the LD within a generation with a fixed scenario, i.e., using fixed TS, in the early stages of a breeding program, to infer across multiple breeding cycles. Although, during breeding cycles (with selection and recombination), LD between markers and QTLs may change due to crossing-over occurrences, especially for more complex traits (Xu, 2013). As a result, there are changes in allelic frequencies and, consequently, allele substitution effects, which may even cause allele fixation (Walsh and Lynch, 2018).

Although previous studies like the ones mentioned above evaluated the effect of markers’ density on genomic prediction models, there are no reports about their influence on a recurrent selection (RS) scheme using models that incorporate additive and dominance effects on cross-pollinated crops. The vast majority of published papers so far have considered only additive models for genomic predictions, which may not represent the genetic complexity of these species. Even though only additive effects are transmitted over generations, the *per se* performance of the population or their single-crosses depends on the dominance deviations (Falconer and Mackay, 1996), as this is the basis of heterosis and crucial in the expression of the agronomic traits of these species (Bernardo, 2010). Additionally, Galli *et al*. (2020) observed negative covariance between single-crosses genetic values predicted from marker effects estimated from parental lines and single-crosses. Hence, this reinforces the inability to predict single-crosses performance based on parental information with additive effects.

Simulation is a suitable tool to verify the effect of marker density reduction in a long-term breeding program using GS. We can control several factors to infer what happens in given genetic parameters over breeding cycles in a fast, cheap and consistent way (Dai *et al*., 2020). It is essential for breeding programs of cross-pollinated crops that usually adopt RS-based methods for obtaining cultivars, either by intrapopulation recurrent selection (IRS) or by reciprocal (RRS), because genetic progress is achieved very slowly with enough selection and recombination over cycles. Additionally, the flexibility of the models and methods used in GS allows integrating allelic interaction effects, as the combining abilities (general and specific) of parentals inbred lines (Werner *et al*., 2018).

Reif *et al*. (2013) observed an increase in the single-crosses performance’s predictive capacity incorporating the model of the effects of the general combining ability (GCA). Despite this, no studies compare the superiority of models that incorporate specific combining ability (SCA) effects concerning genomic best linear unbiased predictor (G-BLUP) model and include additive and dominance effects, mainly if we consider long-term breeding cycles. Thus, our objectives were: (i) to assess the impact of reduction marker density on the prediction accuracy in long-term breeding programs, (ii) to verify how the inclusion of dominance effects has implications on the main genetic parameters, and (iii) to compare genomic prediction models using combining ability, additive and additive – dominance effects over many cycles.

## MATERIAL AND METHODS

### Empirical datasets

#### Phenotypic data

This study considered two maize single-crosses datasets, obtained from Helix Seeds Company (HEL) and the University of São Paulo (USP), phenotyped for the grain yield (GY, in Mg ha^-1^). Plots were manually harvested, and GY was corrected to 13% moisture. Each dataset was analyzed independently. The HEL dataset consists of 452 maize single-crosses obtained by crossing 111 inbred lines. The hybrids were evaluated in a randomized complete block design in five Brazilian sites during the 2015 season. The USP dataset consists of 903 maize single-crosses obtained from a diallel mating design between 49 inbred lines. The single-crosses were evaluated in 2016 and 2017 at two sites (Piracicaba and Anhumas) in São Paulo State, Brazil. Field trials were performed in an augmented block design, each block consisting of 16 unique single-crosses and two commercial hybrids as checks. Single-crosses were evaluated at each site and year under two nitrogen fertilization levels (ideal and low N), composing eight environments. More details about the experimental design, cultivation practices for HEL and USP datasets can be accessed in Sousa *et al*. (2017) and Galli *et al*. (2020), respectively.

#### Genetic-statistical models to estimate BLUEs

Mixed model equations were used for the statistical analysis of the single-crosses at each data set. The joint analysis of each phenotype was performed to estimate genotype means across environments. Then, we fitted the following mixed model to obtain the best linear unbiased estimator (BLUE) for each genotype and then estimate the adjusted means across environments via the *breedR* package (Muñoz and Rodriguez, 2016) in R software (R Development Core Team, 2019). The analysis for the USP dataset were performed by this model:

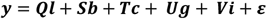

where ***y*** is the vector of phenotypic values of single-crosses and checks; ***l*** is the vector of fixed effects of the environment (combination of site × year × N level); ***b*** is the vector of random effect of block nested within environment, where 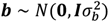 is the vector of fixed effect of checks; ***g*** is the vector of fixed effects of single-crosses; ***i*** is the vector of fixed effects of interaction checks × environments; ***ε*** is the vector of random residuals effects, which are confounded in the final residual term, where ***ε*** ∼ *N*(**0, *D***_***e***_). ***Q, S, T, U***, and ***V*** are the incidence matrices for ***l, b, c, g***, and ***i***. We assumed an unstructured covariance matrix across environments for the residual term (***D***_***e***_).

For phenotypic analysis of the HEL dataset, we use the following reduced model:

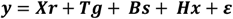

where ***r*** is the vector of block effect considered as random, where 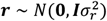 is the vector of random effects of the environment (sites), where 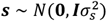 is the vector of the random effects of single-crosses by environment interaction, where 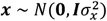; and ***ε*** is the vector of error, where 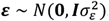. ***X, T, B***, and ***H*** are incidence matrices for ***r, g, s, x***.

Subsequently, for both models, the single-crosses effect was considered random to obtain the variance components, performed by restricted maximum likelihood (REML/BLUP) procedure (Henderson, 1975).

#### Genotypic data

All parental inbred lines (USP and HEL datasets) were genotyped using the Affymetrix^®^ Axiom Maize Genotyping array containing 616 K SNPs (Unterseer *et al*., 2014). As for standard quality controls (QC), markers with low call rate (< 90%) and non-biallelic were removed, and the remaining missing data was imputed by Beagle 5.0 algorithm (Browning et al. 2018). Heterozygous *loci* on at least one individual were removed, and single-crosses genotypes were obtained *in silico* by genomic information from parental inbred lines. Finally, markers with minor allele frequency (MAF) < 0.05 were removed from the hybrid genomic matrix resulting in 63,104 and 30,467 SNP markers for USP and HEL datasets.

#### Markers reduction on datasets

Different SNP marker sets were obtained via two approaches: (i) removing SNP markers based on the LD between them, and (ii) using the genetic algorithm LA-GA-T implemented in the *STPGA R* package that aims to optimize training sets (OTS) developed by Akdemir (2017).

In the first, pairwise LD was calculated as squared allele frequencies correlation (*r*^2^). Then, in order to remove redundant markers, we applied the following threshold values, 0.99, 0.90, 0.80, 070, 0.60, 0.50, 0.40, 0.30, 0.20, 0.10, and 0.01 using the *SNPRelate* package (Zheng et al., 2012). As the benchmark scenario, we used the full matrix of SNP markers (M_Full), after quality control, for downstream comparisons. In the second approach, we applied a singular value decomposition (SVD) using the number of eigenvalues that explained 98% of the variance, according to Pocrnic *et al*. (2016). To reduce computational time and represent only the LD among markers’ physical components, we process SVD + OTS within each chromosome. Lastly, we had thirteen SNP matrices to perform genomic prediction models.

#### Genomic prediction

We used the thirteen SNP matrices to build genomic relationship matrices (GRM). We applied additive and additive-dominance G-BLUP models, respectively, to perform the genomic prediction, following the equations :

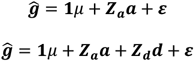

Where ***ĝ*** is the vector of adjusted means of single-crosses from the phenotypic analysis; *μ* is the mean (intercept); ***a*** is the vector of additive genetic effects of the individuals, where 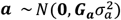;***d*** is the vector of dominance effects, where 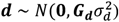; and ***ε*** is the vector of random residuals 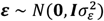. ***1*** is the vector of ones; ***Z***_***a***_ and ***Z***_***d***_ are incidence matrices for ***a*** and 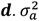 is the genomic additive variance, 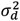 is the genomic dominance variance, and 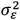 is the residual variance. ***G***_***a***_ and ***G***_***d***_ are the additive and dominance genomic relationship matrices, following the equations: 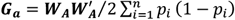 and 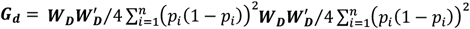, where *p*_*i*_ is the frequency of one allele of the locus *i* and ***W*** is the incidence matrix of markers (VanRaden, 2008). The ***W***_***A***_ matrix was coded as 0 for homozygote A_1_A_1_, 1 for the heterozygote A_1_A_2_, and 2 for the homozygote A_2_A_2_. For ***W***_***D***_, genotypes were coded as 0 for both homozygotes and 1 for heterozygote.

The *snpReady* package (Granato *et al*., 2018) was used to build GRM and obtain estimates of effective population size (N_e_) based on ***G***_***a***_ matrix. ***G***_***a***_ and ***G***_***d***_ matrices obtained from SNP datasets were used to calculate the correlation between them via Mantel correlation using the *vegan* R package (Oksanen *et al*., 2019). The superheat R package built the heatmaps (Barter and Yu, 2018) using M_Full, M_0.5, M_0.01, and M_OTS. For comparison, we used ***G***_***a***_ and ***G***_***d***_ from M_Full dataset to cluster single-crosses using the Euclidean distance. Afterward, the singles-crosses order was used to sort heatmaps M_0.5, M_0.01, and M_OTS. The genomic prediction models were performed using the *sommer* package (Covarrubias-Pazaran, 2016).

Predictive ability was estimated as the correlation between the predicted and observed genotypic values via cross-validation alpha-based design (CV-α, Yassue *et al*., 2020), which is an extension of the methodology presented by Shao (1993) and consists of assigning treatments to folds in each replication using alpha-lattice sorting premises. The CV-α was intended to create scenarios with two, three, or four replicates, regardless of the number of treatments. Each replicate is split into folds, and the number of folds is determined by the percentage of training and validation sets. Each fold across replicates is based on the α(0,1) lattice design aiming to reduce the concurrences of any two treatments in the same fold (block) across the replicates (Patterson and Williams, 1976). We used here four replicates with five-folds, and predictive abilities were averaged.

### Simulated experiments

#### Characteristics of the founder population, trait genetic architecture, and genotyping chips

We simulated a population of maize single-crosses from parental inbred lines to perform phenotypic and genomic predictions for a quantitative trait. For this, we used the *AlphaSimR* package (Gaynor *et al*., 2020), and the scripts can be found in the supplementary files. A historical population founded of 1,000 individuals was simulated stochastically with ten chromosomes containing 31,000 segregating loci. The individuals were diploidized but not inbred.

Trait genetic architecture was simulated considering: (i) only additive effects and (ii) additive + dominance effects. In both, we used randomly sampling 1,000 QTLs (100 per chromosome) with effects drawn from a gamma distribution with a scale parameter of 1.0. We considered an average degree of dominance of 0.5 in each QTL for a trait with additive-dominance effects, according to the ratio between additive effects and dominance in Hallauer *et al*. (2010). We consider heritabilities in the broad and narrow sense equal to 0.5 and 0.33, respectively. From this, it was possible to estimate the error variance component. The phenotypic values were obtained by sum effects of the QTLs plus the random error from the segregating loci with mean 0 and variance 1.

In order to represent the genotyping chips, we used three marker densities sampled from segregating loci: the first was 30K SNPs, which on average is roughly equivalent to 15 markers per cM; the second was 1.5K SNPs equivalent to 1 marker per cM; and the third, 0,75K SNPs means one marker to each 2 cM.

#### Breeding schemes based on recurrent selection

We used two main breeding schemes to evaluate the effect of markers reduction on predictions over 15 cycles: (i) intrapopulation recurrent selection (IRS) and (ii) reciprocal recurrent selection (RRS). To compose the base-population IRS, we selected 15 inbred lines (Fig. 1). Subsequently, a full diallel was composed between them to generate the initial cycle (C_0_). For RRS, the base population was built similarly, but in this case, parents of the best single-crosses were selected to build the two heterotic groups (HG), being ten unique parents in each. All progenies were composed of 50 individuals (equivalent to one trial with 2 plots of 25 plants), regardless of the breeding scheme, stage, or cycle.

**Fig. 1.**
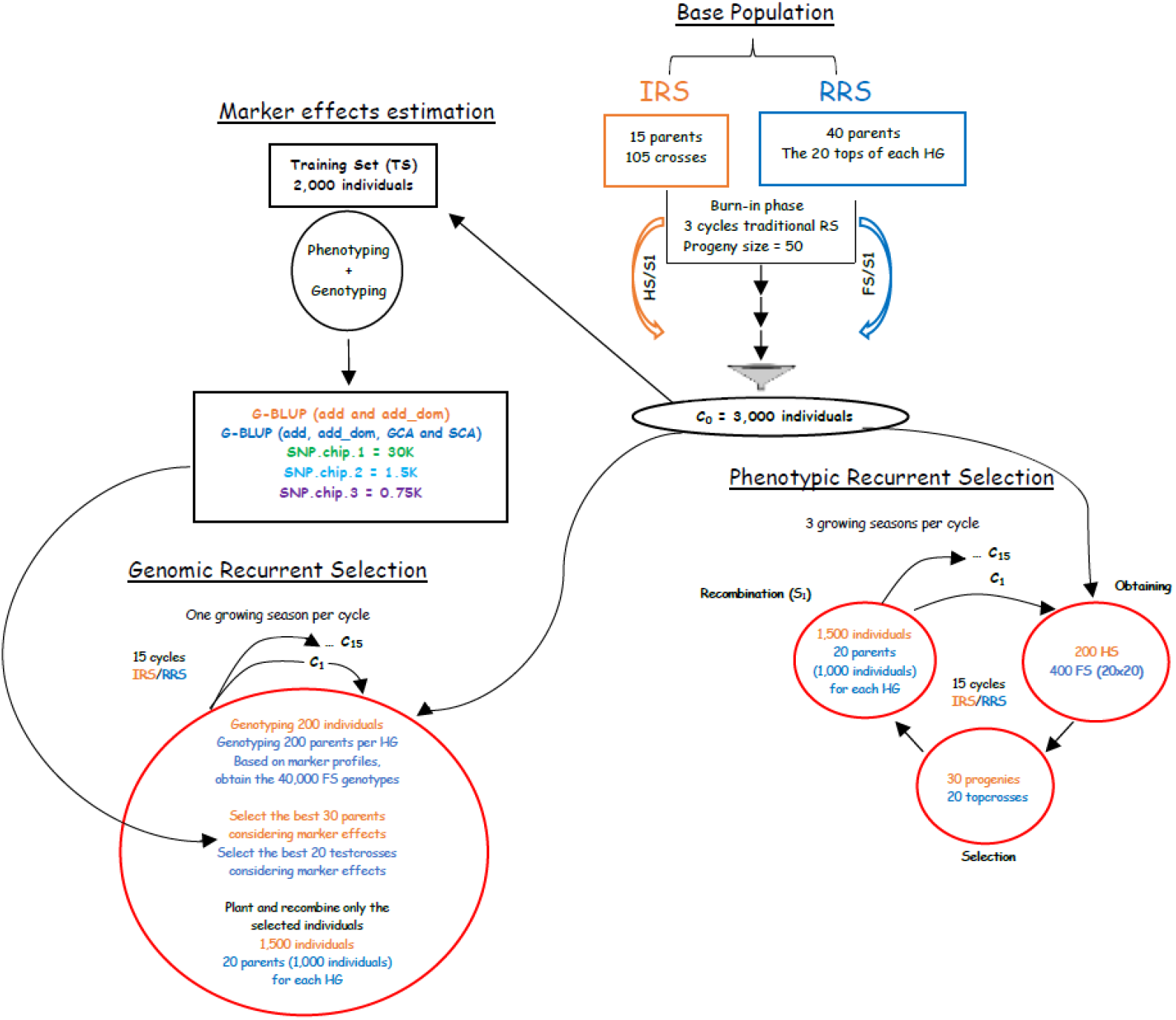
Schematic representation of the breeding schemes based on recurrent selection applied in simulated data. After obtaining the historical population, were developed populations that served as the basis for the IRS (exclusive steps in orange color) and RRS (blue color) schemes. Black information was common to both schemes.

The historical population and trait parameters were equal in both schemes. For the burn-in stage, we used two traditional recurrent selection schemes:

***IRS*** – 200 half-sib progenies were obtained; then, the top 30 progenies were selected, and there was the random recombination of 1,500 self-fertilized individuals (S_1_), corresponding the 30 best parents;

***RRS*** – 200 full-sib progenies were obtained – via North Carolina mating design II (N CII). Then, the best hybrids were selected, and we proceed with the recombination of the 10 best parents for each HG. At the end of each cycle, the fixation index (Fst) was estimated to observe the genetic distance between HG.

After the three cycles of the burn-in phase, 3,000 individuals were randomly selected in each scheme (Fig. 1). Among them, 2,000 were sampled to compose the training set (TS) for genomic predictions, where the marker effects were estimated using the three different SNP-chip densities. For that, genotyping and phenotyping of these individuals were simulated. These 3,000 individuals were also used as the base population (C_0_) for the next 15 recurrent selection cycles (genomic and phenotypic). In each cycle, for both phenotypic breeding schemes, obtaining progenies, selection, and recombination were made based on 200 progenies, similarly to the burn-in phase. Conversely, for the genomic schemes, the genotype of the 200 parents allows us to skip the progeny phase, select directly based on the genomic breeding values, and plant only the select individuals to recombine, closing one breeding cycle in just one growing season. Furthermore, based on the parents’ genotype, we can obtain *in silico* all 40,000 possible single-crosses genotypes (200 x 200), increasing the population size and the intensity of selection.

#### Genomics predictions and comparison models

According to the equations mentioned above, we used additive and additive-dominance G-BLUP models to perform the genomic prediction on three SNP-chip densities in both breeding schemes. Additionally, we model for the RRS scheme with combining abilities effects. In general, the genomic estimated value (GEV) of single-crosses was obtained using the estimates of GCA and SCA according to the following expression:

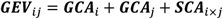

where ***GEV***_ij_ is the genomic estimated value of single-cross *ij*, ***GCA***_*i*_ is the general combining ability of parent *i*, ***GCA***_***j***_ is the general combining ability of parent ***j***, and ***SCA***_*i*×***j***_ is the specific combining ability from the cross between *i* and ***j*** parentals. Kronecker products were used to model the interaction between single-crosses’ parental genomes to capture SCA effects (Acosta-Pech *et al*., 2017; Basnet *et al*., 2019). We repeated the simulations 10 times and averaged the estimates above. Accuracy for phenotypic selection was estimated by the square root of heritability and for GS by the correlation between genetic values and estimated breeding values.

## RESULTS

### Markers reduction on empirical datasets

The number of markers decreased substantially with reducing LD levels on empirical datasets (Table 1). In general, the reduction was smoother in both datasets up to the matrix with 10% LD between markers (M_0.1). Hence, from the matrix with all markers (M_Full) to the one with only 1% LD between markers (M_0.01), we observed a reduction of approximately 11.5 and 3.3 times for the USP and HEL datasets. However, from M_0.01 to matrix markers obtained with the algorithm for the optimization training set (M_OTS), drastic reductions were observed for both datasets, ranging from 3.9 to 5.4 times for USP and HEL.

**Table 1.** Number SNP-markers per chromosome as a function of the marker density reduction by different levels of linkage disequilibrium and OTS algorithm. USP and HEL datasets.

| USP dataset |  |  |  |  |  |  |  |  |  |  |  |  |  |
| --- | --- | --- | --- | --- | --- | --- | --- | --- | --- | --- | --- | --- | --- |
| Chrm/MD | M_FULL | M_0.99 | M_0.9 | M_0.8 | M_0.7 | M_0.6 | M_0.5 | M_0.4 | M_0.3 | M_0.2 | M_0.1 | M_0.01 | M_OTS |
| 1 | 10 689 | 10 548 | 9442 | 8359 | 7504 | 6701 | 5859 | 5049 | 4096 | 3146 | 2152 | 916 | 233 |
| 2 | 8872 | 8789 | 7876 | 7015 | 6296 | 5595 | 4910 | 4175 | 3370 | 2574 | 1675 | 792 | 186 |
| 3 | 5468 | 5420 | 4869 | 4344 | 3897 | 3522 | 3069 | 2614 | 2078 | 1569 | 1057 | 451 | 134 |
| 4 | 5466 | 5394 | 4821 | 4308 | 3873 | 3457 | 3049 | 2604 | 2119 | 1661 | 1159 | 545 | 121 |
| 5 | 8091 | 8007 | 7224 | 6416 | 5794 | 5145 | 4473 | 3802 | 3055 | 2292 | 1515 | 678 | 173 |
| 6 | 5250 | 5197 | 4693 | 4201 | 3783 | 3405 | 2970 | 2554 | 2045 | 1533 | 1036 | 430 | 113 |
| 7 | 6097 | 6045 | 5436 | 4853 | 4397 | 3963 | 3493 | 2956 | 2408 | 1783 | 1197 | 545 | 136 |
| 8 | 6520 | 6440 | 5794 | 5160 | 4650 | 4150 | 3640 | 3093 | 2530 | 1950 | 1247 | 563 | 151 |
| 9 | 3421 | 3379 | 3030 | 2717 | 2464 | 2205 | 1955 | 1665 | 1330 | 1048 | 663 | 303 | 83 |
| 10 | 3230 | 3190 | 2858 | 2584 | 2335 | 2096 | 1843 | 1583 | 1266 | 979 | 652 | 276 | 78 |
| Total | 63 104 | 62 409 | 56 043 | 49 957 | 44 993 | 40 239 | 35 261 | 30 095 | 24 297 | 18 535 | 12 353 | 5499 | 1408 |
| HEL dataset |  |  |  |  |  |  |  |  |  |  |  |  |  |
| 1 | 3977 | 3976 | 3762 | 3558 | 3405 | 3247 | 3059 | 2881 | 2673 | 2442 | 2089 | 1325 | 222 |
| 2 | 3460 | 3460 | 3283 | 3097 | 2948 | 2790 | 2647 | 2487 | 2286 | 2047 | 1736 | 1053 | 188 |
| 3 | 3329 | 3329 | 3170 | 2989 | 2830 | 2685 | 2552 | 2377 | 2241 | 2008 | 1684 | 1045 | 185 |
| 4 | 3057 | 3057 | 2918 | 2771 | 2651 | 2516 | 2389 | 2246 | 2097 | 1880 | 1612 | 1005 | 177 |
| 5 | 3737 | 3734 | 3531 | 3335 | 3179 | 3019 | 2872 | 2703 | 2464 | 2199 | 1832 | 1102 | 212 |
| 6 | 2943 | 2942 | 2766 | 2614 | 2464 | 2332 | 2193 | 2017 | 1854 | 1614 | 1327 | 803 | 172 |
| 7 | 3039 | 3037 | 2856 | 2695 | 2551 | 2418 | 2286 | 2147 | 1977 | 1765 | 1428 | 860 | 173 |
| 8 | 2615 | 2613 | 2452 | 2328 | 2230 | 2125 | 2010 | 1872 | 1718 | 1523 | 1299 | 812 | 147 |
| 9 | 2242 | 2241 | 2133 | 2018 | 1935 | 1844 | 1746 | 1648 | 1524 | 1379 | 1132 | 705 | 138 |
| 10 | 2068 | 2067 | 1976 | 1875 | 1790 | 1709 | 1628 | 1515 | 1388 | 1234 | 1021 | 623 | 120 |
| Total | 30 467 | 30 456 | 28 847 | 27 280 | 25 983 | 24 685 | 23 382 | 21 893 | 20 222 | 18 091 | 15 160 | 9333 | 1734 |

Although considerable markers reduction over LD levels, the effective population size (Ne) has not been changed for both single-crosses datasets when considering the different matrices (scenarios) for estimating this parameter (Table 2). In addition, the significant and high correlation coefficients between additive (***G***_***a***_) (> 0.98 for USP and HEL datasets) and dominance (***G***_***d***_) matrices (> 0.73 for USP and > 0.99 for HEL dataset) obtained from different markers densities (Tables S1 and S2), reiterate the matrices with low-density markers are as representative as those with high-density. Besides that, heatmaps obtained with additive and dominance matrices among the most extreme scenarios prove that the genetic relationship has not been altered with the marker density reduction (Figs. S1 and S2).

**Table 2.** Effective population size (Ne) obtained by additive matrices (*A*) with markers density reduction in different linkage disequilibrium levels and the OTS algorithm. USP and HEL datasets.

| MD | Ne |  |
| --- | --- | --- |
|  | USP | HEL |
| M_FULL | 456.21 | 230.55 |
| M_0.99 | 456.19 | 230.55 |
| M_0.9 | 456.26 | 230.48 |
| M_0.8 | 456.37 | 230.43 |
| M_0.7 | 456.32 | 230.38 |
| M_0.6 | 456.33 | 230.26 |
| M_0.5 | 456.30 | 230.26 |
| M_0.4 | 456.55 | 230.14 |
| M_0.3 | 456.59 | 230.12 |
| M_0.2 | 456.73 | 230.10 |
| M_0.1 | 456.85 | 230.16 |
| M_0.01 | 457.56 | 231.33 |
| M_OTs | 455.88 | 229.84 |

Similar to the previous results for the USP and HEL datasets, we noted that the predictive ability has remained mostly unchanged (minor changes) throughout the various markers reduction scenarios, including the most extremes ones (Fig. 2). Therefore, with approximately 1,500 markers (1,408 for USP and 1734 for HEL datasets), it is possible to predict maize single-crosses’ performance with the same reliability as all marks after the quality control process. Additionally, greater predictive abilities were observed when we adopted the additive-dominance model regardless of marker density. On average, this model surpassed the compound only by additive effects by 9.8% and 19% for the USP and HEL datasets, respectively.

**Fig. 2.**
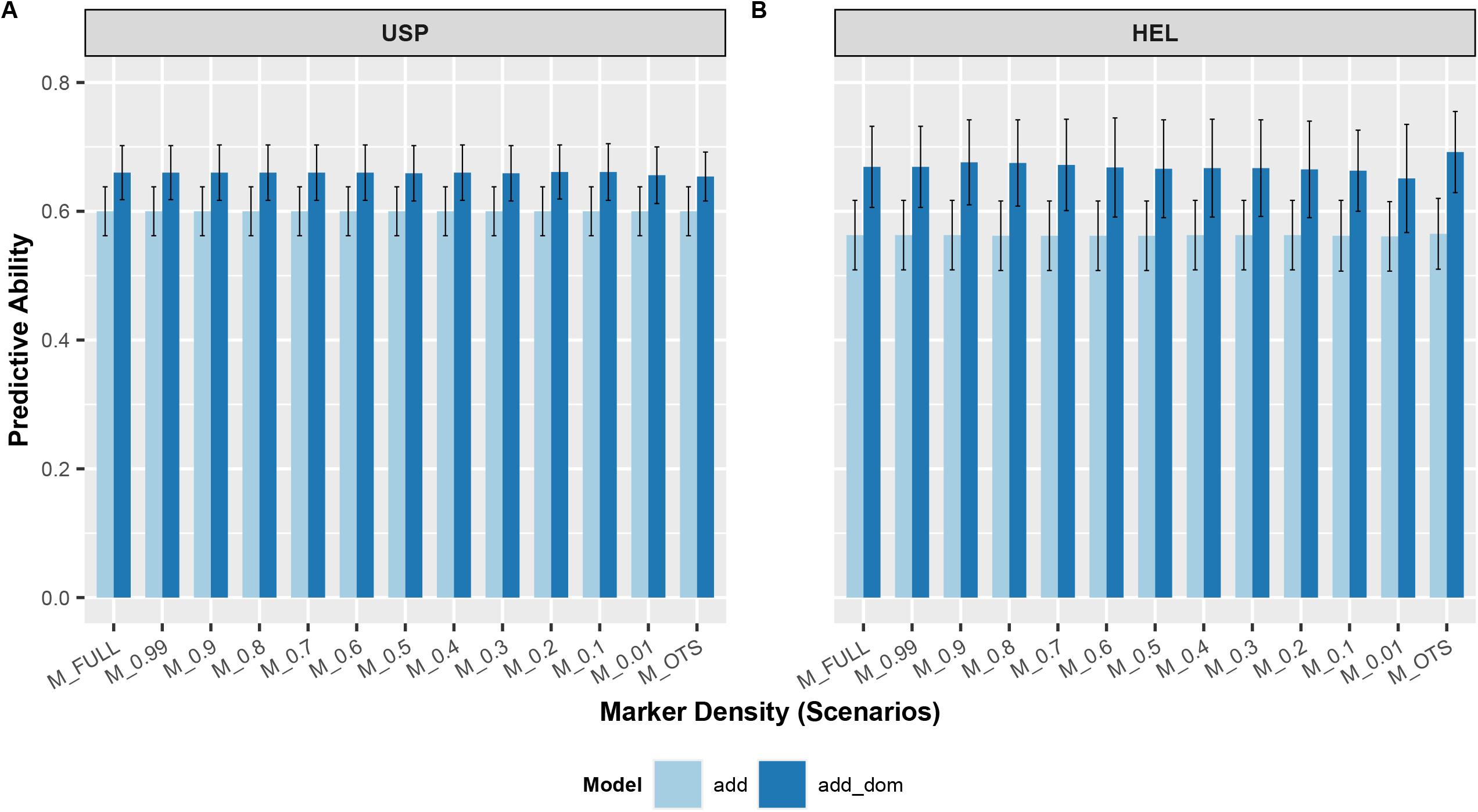
Predictive ability with several scenarios of markers density reduction by different levels of linkage disequilibrium and algorithm for OTS using additive and additive-dominant models. **A** USP dataset **B** HEL dataset.

### Marker density reduction effect on long-term IRS breeding

The simulated data showed that GS overcame phenotypic selection for the population mean during all breeding cycles in an IRS scheme, regardless of marker density (Fig. 3A). GS’s superiority began to become evident in the third cycle. At the end of the 15 cycles, the GS allowed an average of 14.5 Mg ha^-1^ to be reached, while the phenotypic selection a maximum of 88% of this. We also observed that for this parameter, the additive model tended to be superior to the additive-dominance, regardless of the selection method (phenotypic or genomic). However, in the phenotypic selection, the difference between models tended to be more significant over the cycles.

**Fig. 3.**
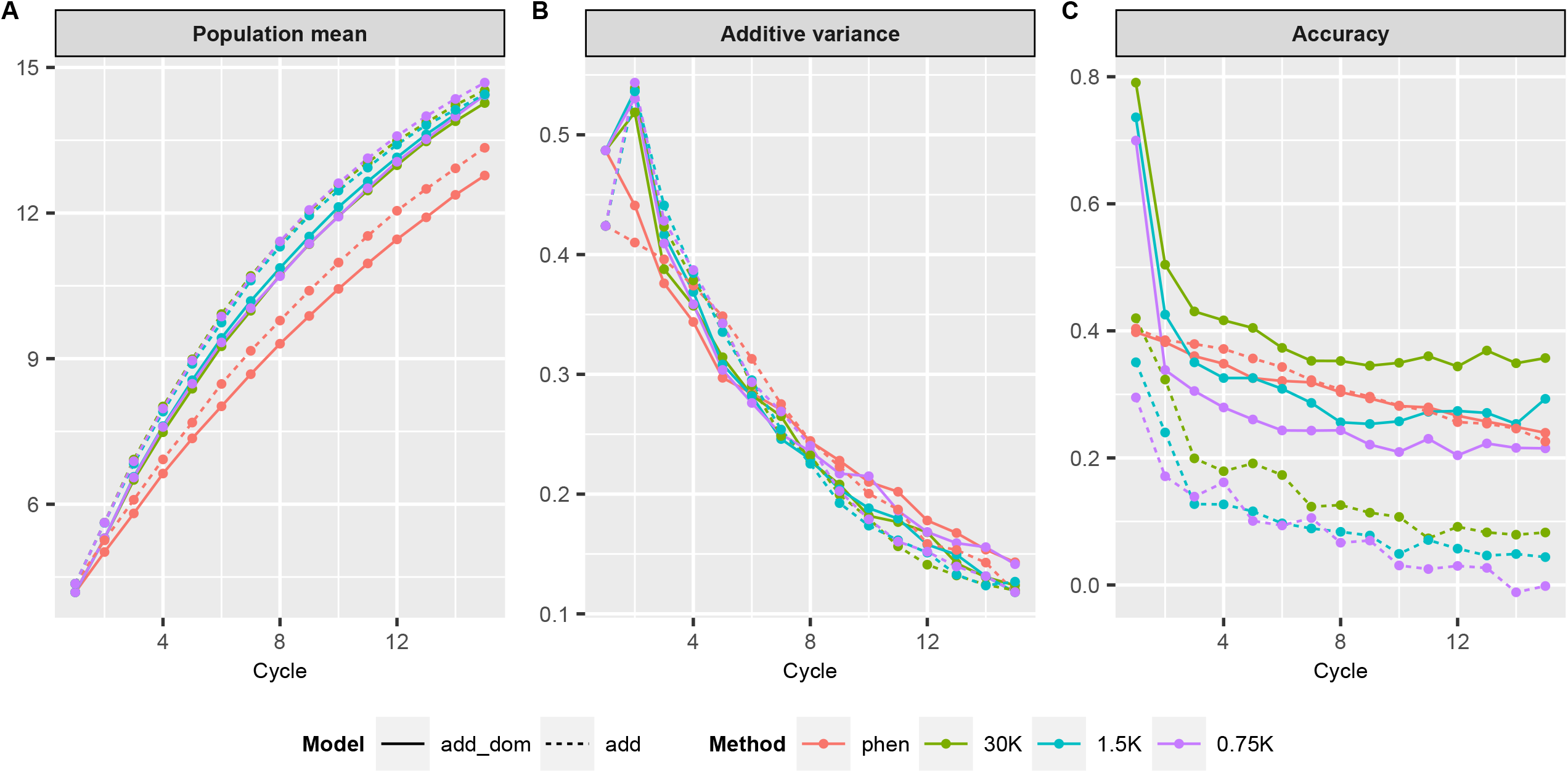
Simulation of 15 breeding cycles via phenotypic and genomic selection (with three marker densities) by the recurrent intrapopulation selection (IRS) scheme with additive and additive-dominant models. **A** Population mean **B** Additive variance **C** Accuracy

The additive variance available for GS increased until the second breeding cycle and was consumed similarly to the phenotypic selection (Fig. 3B). Selection methods and genetic models did not show a significant difference of additive variance tendencies over breeding cycles. On the other hand, GS using the additive-dominance model generated higher accuracy estimates (Fig. 3C). In the initial cycle, GS with additive-dominance model reached almost twice the accuracy (> 0.7) of GS with the additive model (maximum 0.43) and also from phenotypic selection regardless of the model (at most 0.40). In general, after the second cycle, GS showed abrupt reductions in reliability. As for the phenotypic selection, there was a loss in accuracy over the cycles, but with a smaller variation. However, higher accuracy was obtained when the breeding program was conducted with GS, and more genomic information was available, regardless of the genetic model. It is worth noting that after the reduction in accuracy of the second cycle, GS using the SNP-chip of 30K promoted less variation and a superior accuracy over other scenarios. Furthermore, the additive-dominance model outperformed the additive one in terms of GS accuracy for IRS long-term breeding.

### Marker density reduction effect on long-term RRS breeding

When the RRS scheme conducts the breeding program, similar outcomes were observed for the population mean (Fig. 4A). Also, the differences between this scheme’s methods were more significant, with performance above 15 Mg ha^-1^ for GS and around 9 Mg ha^-1^ for phenotypic selection at the end of the 15 cycles. The best crosses obtained via GS also outperform those obtained via phenotypic selection, regardless of SNP-chip densities (Fig. 4B). In general, single-crosses with the adoption of GS exceeding 17 Mg ha^-1^, on average 70% higher than those obtained by phenotypic selection. GS allowed maintaining greater genetic divergence by fixation index (Fst) between heterotic groups (Fig. 4C), especially from the eighth cycle.

**Fig. 4.**
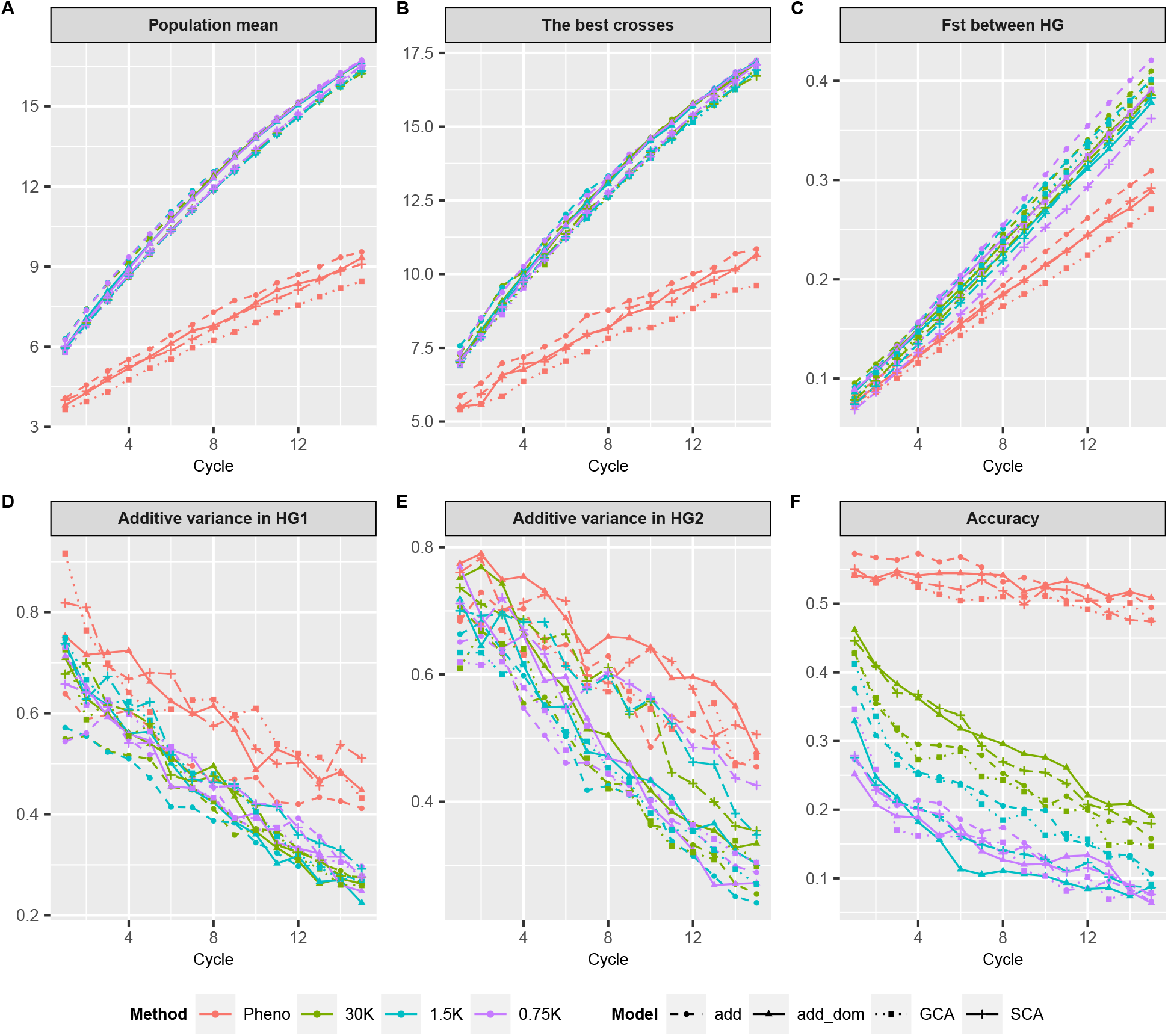
Simulation of 15 breeding cycles via phenotypic and genomic selection (with three marker densities) by the reciprocal recurrent selection (RRS) scheme with additive, additive-dominant, CGA, and SCA G-BLUP models. **A** Population mean **B** Best crosses **C** Fst between HG **D** Additive variance HG1 **E** Additive variance HG 2 **F** Accuracy

There was a similar behavior for the population mean and best crosses regardless of the models’ GS method. However, for the Fst parameter, the additive model showed a slightly higher tendency than the additive-dominance model. The phenotypic selection follows the fashion showing the additive model’s superiority for the population mean and best crosses parameters. On the other hand, the GCA model tended to be inferior in all of them. For population mean, the best-crosses, and Fst, the additive-dominance and SCA models presented practically the same performance. This same behavior was observed when we considered the phenotypic selection for the best crosses and population mean parameters, as seen in the IRS scheme.

Unlike what was observed in the IRS scheme, the additive variance (for both heterotic groups) tended to be consumed more quickly when GS conducted the breeding program, regardless of the marker density and model (Fig. 4D and 4E). The phenotypic selection also showed greater accuracy from the beginning to the end of breeding cycles. (Fig. 4F). Overall, the accuracy tended to be superior in the breeding process with a higher density of markers. The modeling did not interfere decisively in the performance of this parameter. However, there was a slight tendency to superiority the additive-dominant model for accuracy, similar to the IRS scheme.

## DISCUSSION

The high-density genotyping chips permit whole coverage of the genome with markers, making it possible to capture the total genetic variability of the trait across the genome and predict the breeding value without collecting phenotypic data of the individuals (Meuwissen *et al*., 2001).

Low-density marker chips are desirable to implement GS in breeding programs to reduce genotyping costs, especially for species with low profitability. Several studies have documented the benefits of using marker subsets in GS analyses (Gorjanc *et al*., 2015; Tayeh *et al*., 2015; Ma *et al*., 2016; Li *et al*., 2018; Sousa *et al*., 2019). Thus, the fact that few markers are sufficient to generate predictive ability similar to those obtained with high markers density, without changing the genetic relationship (Tables 1 and 2; Figs. S1 and S2) brings excellent advantages for the low-density chips, leading to technology improvements and cost (Heaton *et al*., 2005; Chessa *et al*., 2007), as well as computational demand.

Nevertheless, long-term recurrent selection difficulties GS effectiveness. Over cycles, recombination between markers and QTL will cause LD decreases, while selection and drift will potentially act to generate new LD or tighten the LD between closely linked *loci* (Hill and Robertson, 1968; Lorenz *et al*., 2011). Thus, there is a demand to study the effects of reducing the markers density over breeding cycles. Therefore, simulations of different scenarios and controlled variants may do this quickly and less costly.

### Consequences of marker reduction for GS to long-term breeding schemes

#### Breeding program conducted by IRS scheme

When the breeding program is conducted by recurrent selection with a single population (IRS), generally, the purpose is to release open-pollinated varieties. Thus, the population means is an essential parameter to evaluate the performance of the IRS scheme. Our results showed that GS applied in an IRS scheme overcomes phenotypic selection, regardless of SNP-chip density (Fig. 3A). This was expected because GS is more informative than phenotypic selection. With the availability of genomic and phenotypic data, it becomes easier to identify superior genotypes with precision. Despite this, the additive variance was consumed by both strategies (phenotypic and genomic selection) similarly (Fig. 3B).

The GS accuracy decreased abruptly from the second breeding cycle (Fig. 3C). That suggests that TS must be re-calibrated (updated) during recurrent selection to maintain prediction ability (Neyhart *et al*., 2017). Indeed, studies simulating recurrent selection in eucalyptus (Jannink, 2010), barley (Denis and Bouvet, 2013), and with empirical data from advanced-cycle rye (Auinger *et al*., 2016) revealed that the accuracy of GS was improved by recalibration of the TS. Therefore, each recurring selection cycle’s recombination events should shift in the QTL-marker LD pattern, making its capture unfeasible. Further, the change in allele frequencies due to the selection over the cycle’s impact on estimating the effects of allelic substitution and, consequently, on the genomic prediction.

Furthermore, this situation can be aggravated when the markers density is initially very low, and then it will compromise the recalibration of the TS. Our results showed a lower loss of accuracy over the breeding cycle when higher marker densities were used (Fig. 3C). Müller *et al*. (2017) illustrated by simulations that more remarkable persistence of predictive accuracy and, consequently, selection gains when higher markers density were used. Thus, most markers seem to buffer better the effect of the loss LD between markers and QTLs over the recombination cycles, even without TS recalibration. Although there is an expectation with reducing genotyping costs with low-density chips, in practice, budgets in companies specialized in genotyping show that the price differences between chips of high and low markers density are negligible (personal communication). Therefore, our results showed that genotyping with low-density chips for a breeding program conducted to long-term breeding cycles is not compensatory.

#### Breeding program conducted by RRS scheme

In RRS, two populations are improved simultaneously, not to increase their performance by themselves but the cross between them. In principle, the frequencies desirable alleles in each population are maximized and improved of additive effects. Subsequently, both populations are crossed to capitalize on heterozygous individuals’ non-additive effects (Hallauer *et al*. 2010). Thus, the RRS is more complex than the IRS. However, when conducted with the GS support, it is possible to run similarly to the IRS scheme. The population performance was better (Fig. 3A and 4A) probably because single-crosses are generated (with more significant heterosis than the open-pollinated varieties).

The performance of single-crosse selected by GS was higher than phenotypic selection (Figs. 4A and 4B). It may be explained due to GS permits to generate *in-silico* all single-crosses combinations (in the present study 40,000), which is unable with traditional phenotypic selection, which increases the chances of identifying the best crosses. Furthermore, the genomic information allows discriminating alleles belonging to each heterotic group and keeping them more divergent over the breeding cycles (Fig. 4C). As the selection is made based on single-crosses, which have a high heterosis effect, the GS better captures the complementarity, heterozygosity, and genetic distance components. That is important for a breeding program that aims to release single-crosses since those are the main components of heterosis (Moll *et al*., 1965; Prasad and Singh, 1986).

Long-term recurrent selection may reduce genetic variance to the point where genetic gains are limited (Jannink, 2010). Indeed, genetic variance showed a substantial decline over breeding cycles in both selection schemes and methods. However, in the RRS scheme, GS resulted in a higher genetic diversity loss than phenotypic selection (Figs. 4D and 4E). Muleta *et al*. (2019) observed similar results with higher losses in the simulation of an oligogenic trait with high heritability and lower for a polygenic trait with low heritability. These authors attributed this fact to higher selection intensity in GS due to more selection cycles per year (simulated two and three cycles of GS per year). In our study, we simulate only one cycle per year for both GS and phenotypic selection. Even so, it was clear that the additive variance is consumed more quickly by GS. Essentially, selection intensity (standardized) will be higher in GS compared to phenotypic selection because it has a larger sample, as previously mentioned.

Maintenance of prediction accuracy across selection cycles is critical for long-term genetic gain (Müller *et al*., 2017). Interestingly, the phenotypic selection also overcame the GS for accuracy throughout cycles (Fig. 4F). Mathematically, the accuracy is a function of the additive variance. Therefore, a reduction of that will promotes a decrease in prediction accuracy. This finding may be explained by the breakdown of LD between markers and QTL due to recombination (Jannink, 2010). Moreover, over cycles to selection and drift of alleles are not tagged by markers in TS, which cannot be targeted by GS (Rutkoski *et al*., 2015), culminating in low accuracy. Another suggested explanation for this result is that the selection is based on genetic divergence, in which the alleles present in individuals captured from both heterotic groups change in direction and sense (upper and lower tail), affecting the reliability of the GS.

Similarly to the IRS scheme, we noted that the higher marker density results in a higher accuracy over the breeding cycles. A larger marker quantity should better capture the LD between QTLs and markers since some of them can disassociate from the QTLs due to recombination during the cycles, whereas other markers can remain associated with the QTLs, maintaining some level of LD.

#### Comparison between prediction models

The predictions by GS tended to be more accurate than phenotypic selection because it considers the real genetic relatedness between genotypes rather than the average expectation. Studies have been conducted with empirical data (Al-Tobasei *et al*., 2020) and with simulations (Hou *et al*., 2020) whether for animal or plant breeding, to understand the markers’ effect on the accuracy of genomic predictions. However, they only consider the additive genetic effects to identify marker subsets and, consequently, make the predictions.

Nevertheless, several other studies have already shown that information about the dominance deviations incorporated via kernels are essential to increase the efficiency of the prediction models GS (Azevedo *et al*., 2015; Dias *et al*., 2018; Matias *et al*., 2018; Dai *et al*., 2020), especially when the species is cross-pollination and heterosis is explored in the final product (Santos *et al*., 2016). Our reports for empirical datasets (Fig. 2) corroborate these others that showed the importance of incorporating additive effects and deviations of dominance in the model to increase predictive ability (Technow *et al*., 2012; Alves *et al*., 2019). In the presence of dominance, the additive-dominant model is expected to be more accurate because the additive model falsely assumes that residuals are independent and identically distributed (Duenk *et al*., 2017). Furthermore, in addition, the additive-dominant models allow separating the effects better than the models that consider only the additive effects. When we consider only the additive kernel in the models, there is a “confusion” among additive effects, dominance, and residuals (Alves *et al*., 2019).

However, in the long-term breeding, we found that the additive-dominance model was consistently superior to the IRS scheme’s accuracy parameter. On the other hand, the additive model tended to be slightly superior to the others for population means (for both IRS and RRS), the best crosses, and Fst between heterotic groups. Duenk *et al*. (2017) showed that the additive model contributed to the error with sampling deviations of genotype frequencies from Hardy-Weinberg Equilibrium (HWE). Conversely, the additive-dominant model is only sampling deviations of allele frequencies. Consequently, according to these authors, in the presence of dominance, the root mean squared error of the average effect of allelic substitution with the additive-dominant model was always smaller, especially when the heritability is low. Therefore, for cross-pollinated species, in which single-cross are commercially exploited, it is expected genetic models that include dominance more robust and generated higher accuracies of the average effect of allelic substitution than purely additive models when dominance is present. However, our results show that this benefit did not have advantages in estimating the parameters studied.

It is expected that GS does not consistently adjust to RRS, since in the modeling process, only a vector of additive effects and deviations dominance is considered for the entire population and their respective heterotic groups. Thus, we use two more approaches in the simulation of breeding cycles with RRS; G-BLUP modeled only with the GCA effects and the other with SCA effects. Based on quantitative genetics theory, it is expected that via traditional GS, individuals from both heterotic groups would be selected by the same allele effects, which might lead to a reduction in the genetic distance among HG over the cycles, vanishing the heterosis expression (Hallauer *et al*., 2010).

In theory, it is hoped that breeders who exploit heterosis to generate products (especially single-crosses) can benefit from models that can predict non-additive genetic effects. Thus, it is possible to predict the performance of single-crosses with greater accuracy. In essence, the two models that best capture dominance deviations are the additive-dominance and SCA. These models presented similar estimates for the parameters studied, such as, for example, population means, best crosses, and Fst. Interestingly, Werner *et al*. (2018) used different models to verify the increase in predictive ability in oilseed rape with the GS approach. This study found that incorporating SCA effects combined with GCA into the RR-BLUP method (similar to G-BLUP) was the approach that caused the most fluctuation over 200 validation cycles. Except for this, all methods (including those based on Bayesian statistics) revealed comparatively robust predictive ability and high accuracy. However, we did not notice any trends of considerable differences between results generated by four models over the 15 cycles, although it is known that the dominance deviations captured by the SCA effect are paramount for heterosis (Falconer and Mackay, 1996).

We believe that the similarity of results is because we select the best single-crosses first, then recombine their parents. This procedure is standard in the RRS scheme and does not change with both modeling approaches. Regarding the single-cross genomic prediction, it is well-known that the non-additive effects are partially included in the parents breeding value, GCA, or in allele substitution effects (Falconer and Mackay, 1996; Bernardo, 2010; Werner *et al*., 2018). Even though it is noteworthy that even in some circumstances generating lower estimates than the additive, the additive-dominance model generates more reliable estimates (Alves *et al*., 2019). Reif *et al*. (2007) demonstrated that the dominance variance decreases concerning additive variance but increases according to the populations’ divergence. The dominance effects are increasingly absorbed by the population mean or become inseparable from additive effects over breeding cycles (Technow *et al*., 2014).

#### Final considerations

It is challenging to disentangle the different factors that may contribute to heterogeneity and predictive ability failure in real data sets (Dai *et al*., 2020). However, several genetic-statistical models and tools are available to minimize bias. Using simulations, we can control several uncontrolled factors in the field and focus on answer specific questions. Our results clearly showed that reducing the markers is punctually efficient (within a breeding cycle or a static group of parents). Hence, the low-density marker sets are not affordable for recurrent genomic selection (either IRS or RRS), usually conducted for the long-term. Overall, the more markers, the higher the genotyping chip’s buffering capacity to support LD losses (between markers and QTL) by crossing-over during recombination cycles. Therefore, this study complements the results of others published recently. Although GS allows a higher genetic gain, especially when weighted by time, Muleta *et al*. (2019) also observed higher GS accuracy losses about phenotypic selection throughout the breeding cycles. In this sense, it is evident the need for more studies about how often and how to make updates to the TS to maximize the reliability and prediction accuracy (Rincent *et al*., 2012), keeping the LD between markers and QTLs (Neyhart *et al*., 2017), and the relationship between TS and target populations satisfactory levels. Finally, we recommend that genotyping should be done with a high marker density so that this in a long-term breeding program maintains the ability to capture LD over cycles and does not become a bottleneck. The additive model is sufficient to predict breeding populations’ perf ormance, but the additive-dominance generates slightly better estimates of accuracy throughout the breeding cycles. Furthermore, regarding non-additive genetic effects, there are no differences between the additive-dominance and SCA G-BLUP models.

## Acknowledgments

We thank Luiz de Queiroz College of Agriculture (University of São Paulo), Federal University of Ceará and Conselho Nacional de Desenvolvimento Científico e Tecnológico (CNPq) – Process 104371/2019-6, for the financial support.

## Data archiving

https://data.mendeley.com/datasets/4nccgtcpgn/1

