## Supplementary material for "Reduction of genotyping marker density for genomic selection is not an affordable approach to long-term breeding in cross-pollinated crops"

**Table S1 Coefficients of correlations between matrices obtained with markers density reduction by different linkage disequilibrium levels and algorithm for OTS**. On the upper diagonal, the additive kinships, and the lower diagonal, the dominance deviations kernels. USP datasets

| | M_FULL | M_0.99 | M_0.9 | M_0.8 | M_0.7 | M_0.6 | M_0.5 | M_0.4 | M_0.3 | M_0.2 | M_0.1 | M_0.01 | M_OTS |
|---|---|---|---|---|---|---|---|---|---|---|---|---|---|
| **M_FULL** | | 1.000 | 1.000 | 1.000 | 1.000 | 1.000 | 0.999 | 0.999 | 0.999 | 0.999 | 0.998 | 0.996 | 0.994 |
| **M_0.99** | 1.000 | | 1.000 | 1.000 | 1.000 | 1.000 | 1.000 | 0.999 | 0.999 | 0.999 | 0.998 | 0.996 | 0.994 |
| **M_0.9** | 0.998 | 0.998 | | 1.000 | 1.000 | 1.000 | 1.000 | 1.000 | 0.999 | 0.999 | 0.998 | 0.996 | 0.994 |
| **M_0.8** | 0.997 | 0.997 | 0.999 | | 1.000 | 1.000 | 1.000 | 1.000 | 0.999 | 0.999 | 0.998 | 0.996 | 0.993 |
| **M_0.7** | 0.995 | 0.995 | 0.997 | 0.998 | | 1.000 | 1.000 | 1.000 | 1.000 | 0.999 | 0.998 | 0.996 | 0.993 |
| **M_0.6** | 0.993 | 0.993 | 0.995 | 0.996 | 0.998 | | 1.000 | 1.000 | 1.000 | 0.999 | 0.998 | 0.996 | 0.993 |
| **M_0.5** | 0.990 | 0.991 | 0.993 | 0.994 | 0.995 | 0.997 | | 1.000 | 1.000 | 0.999 | 0.999 | 0.996 | 0.993 |
| **M_0.4** | 0.987 | 0.987 | 0.990 | 0.991 | 0.992 | 0.993 | 0.995 | | 1.000 | 0.999 | 0.999 | 0.996 | 0.992 |
| **M_0.3** | 0.982 | 0.982 | 0.984 | 0.985 | 0.986 | 0.987 | 0.988 | 0.991 | | 0.999 | 0.999 | 0.996 | 0.992 |
| **M_0.2** | 0.973 | 0.973 | 0.975 | 0.976 | 0.977 | 0.978 | 0.978 | 0.979 | 0.981 | | 0.999 | 0.996 | 0.991 |
| **M_0.1** | 0.954 | 0.954 | 0.955 | 0.955 | 0.956 | 0.957 | 0.957 | 0.957 | 0.958 | 0.959 | | 0.997 | 0.990 |
| **M_0.01** | 0.730 | 0.895 | 0.894 | 0.894 | 0.894 | 0.893 | 0.893 | 0.893 | 0.893 | 0.892 | 0.893 | | 0.987 |
| **M_OTS** | 0.823 | 0.822 | 0.821 | 0.819 | 0.818 | 0.816 | 0.814 | 0.810 | 0.806 | 0.797 | 0.779 | 0.895 | |

All correlation coefficients involving pairs of information were significant at p <0.01 by the Mantel test.

**Table S2 Coefficients of correlations between matrices obtained with markers density reduction by different linkage disequilibrium levels and algorithm for OTS**. On the upper diagonal, the additive kinships, and the lower diagonal, the dominance deviations kernels. HEL datasets

| | **M_FULL** | **M_0.99** | **M_0.9** | **M_0.8** | **M_0.7** | **M_0.6** | **M_0.5** | **M_0.4** | **M_0.3** | **M_0.2** | **M_0.1** | **M_0.01** | **M_OTS** |
|---|---|---|---|---|---|---|---|---|---|---|---|---|---|
| **M_FULL** | | 1.000 | 1.000 | 1.000 | 1.000 | 1.000 | 1.000 | 0.999 | 0.999 | 0.999 | 0.999 | 0.997 | 0.993 |
| **M_0.99** | 1.000 | | 1.000 | 1.000 | 1.000 | 1.000 | 1.000 | 0.999 | 0.999 | 0.999 | 0.999 | 0.997 | 0.993 |
| **M_0.9** | 0.996 | 1.000 | | 1.000 | 1.000 | 1.000 | 1.000 | 1.000 | 0.999 | 0.999 | 0.999 | 0.997 | 0.993 |
| **M_0.8** | 0.996 | 1.000 | 0.993 | | 1.000 | 1.000 | 1.000 | 1.000 | 1.000 | 0.999 | 0.999 | 0.996 | 0.993 |
| **M_0.7** | 0.996 | 1.000 | 1.000 | 0.993 | | 1.000 | 1.000 | 1.000 | 1.000 | 0.999 | 0.999 | 0.996 | 0.993 |
| **M_0.6** | 0.996 | 1.000 | 1.000 | 1.000 | 0.993 | | 1.000 | 1.000 | 1.000 | 1.000 | 0.999 | 0.996 | 0.993 |
| **M_0.5** | 0.996 | 0.999 | 1.000 | 1.000 | 1.000 | 0.993 | | 1.000 | 1.000 | 1.000 | 0.999 | 0.996 | 0.993 |
| **M_0.4** | 0.996 | 0.999 | 1.000 | 1.000 | 1.000 | 1.000 | 0.993 | | 1.000 | 1.000 | 0.999 | 0.996 | 0.993 |
| **M_0.3** | 0.996 | 0.999 | 0.999 | 1.000 | 1.000 | 1.000 | 1.000 | 0.992 | | 1.000 | 0.999 | 0.996 | 0.993 |
| **M_0.2** | 0.996 | 0.999 | 0.999 | 1.000 | 1.000 | 1.000 | 1.000 | 1.000 | 0.992 | | 1.000 | 0.997 | 0.992 |
| **M_0.1** | 0.997 | 0.999 | 0.999 | 0.999 | 0.999 | 1.000 | 1.000 | 1.000 | 1.000 | 0.992 | | 0.997 | 0.992 |
| **M_0.01** | 0.991 | 0.996 | 0.999 | 0.999 | 0.999 | 0.999 | 0.999 | 0.999 | 0.999 | 1.000 | 0.992 | | 0.988 |
| **M_OTS** | 0.993 | 0.993 | 1.000 | 1.000 | 1.000 | 1.000 | 0.999 | 0.999 | 0.999 | 0.999 | 0.999 | 0.996 | |

All correlation coefficients involving pairs of information were significant at $p < 0.01$ by the Mantel test.

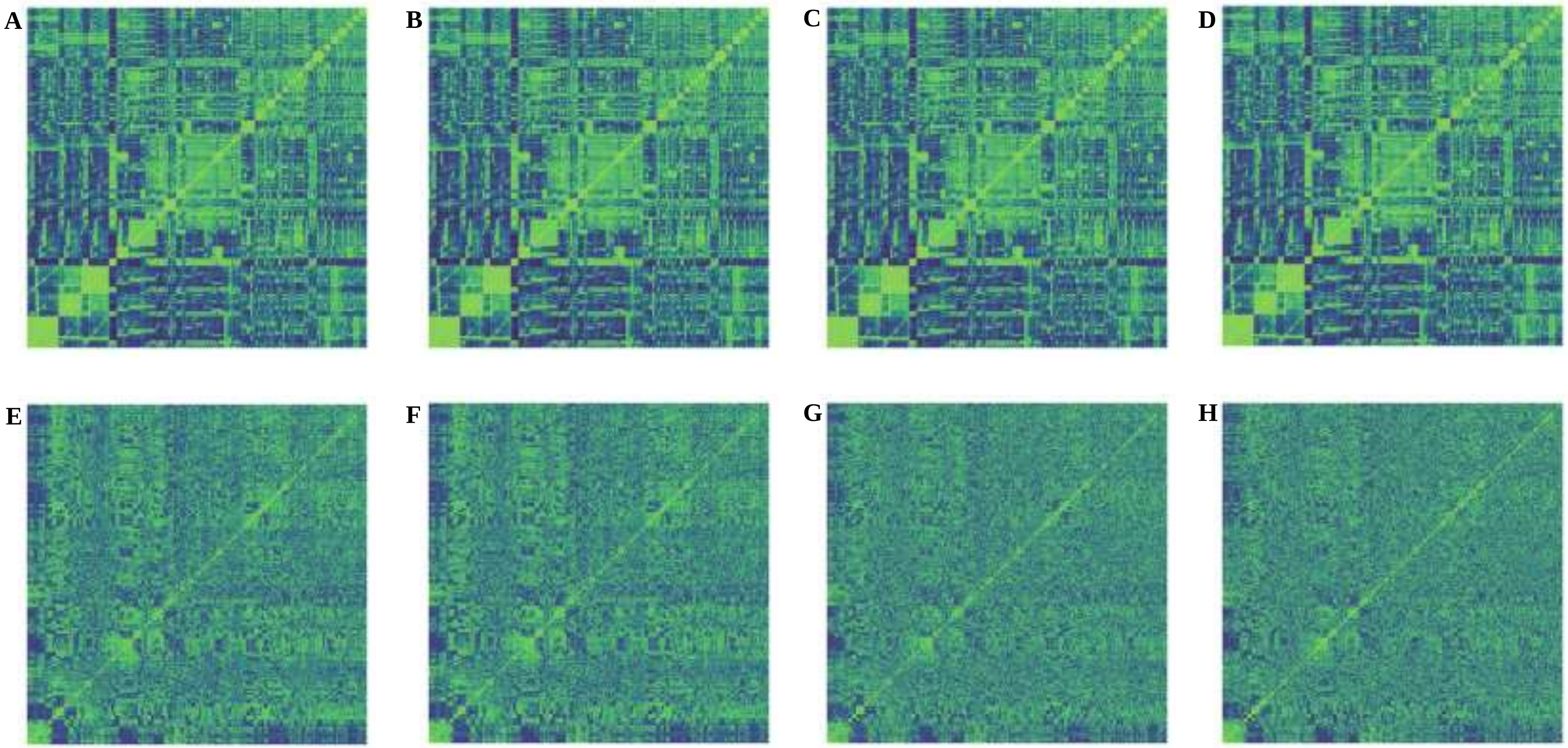


**Fig. S1 Some heatmaps for the additive (figures at the top) and dominance (figures at the bottom) kinships. A and E** Full-matrix (M_full) **B and F** reduced by 0.5 LD (M_0.5) **C and G** by 0.01 LD (M_0.01) and **D and H** optimized via algorithm for OTS (M_OTS), respectively. USP dataset.

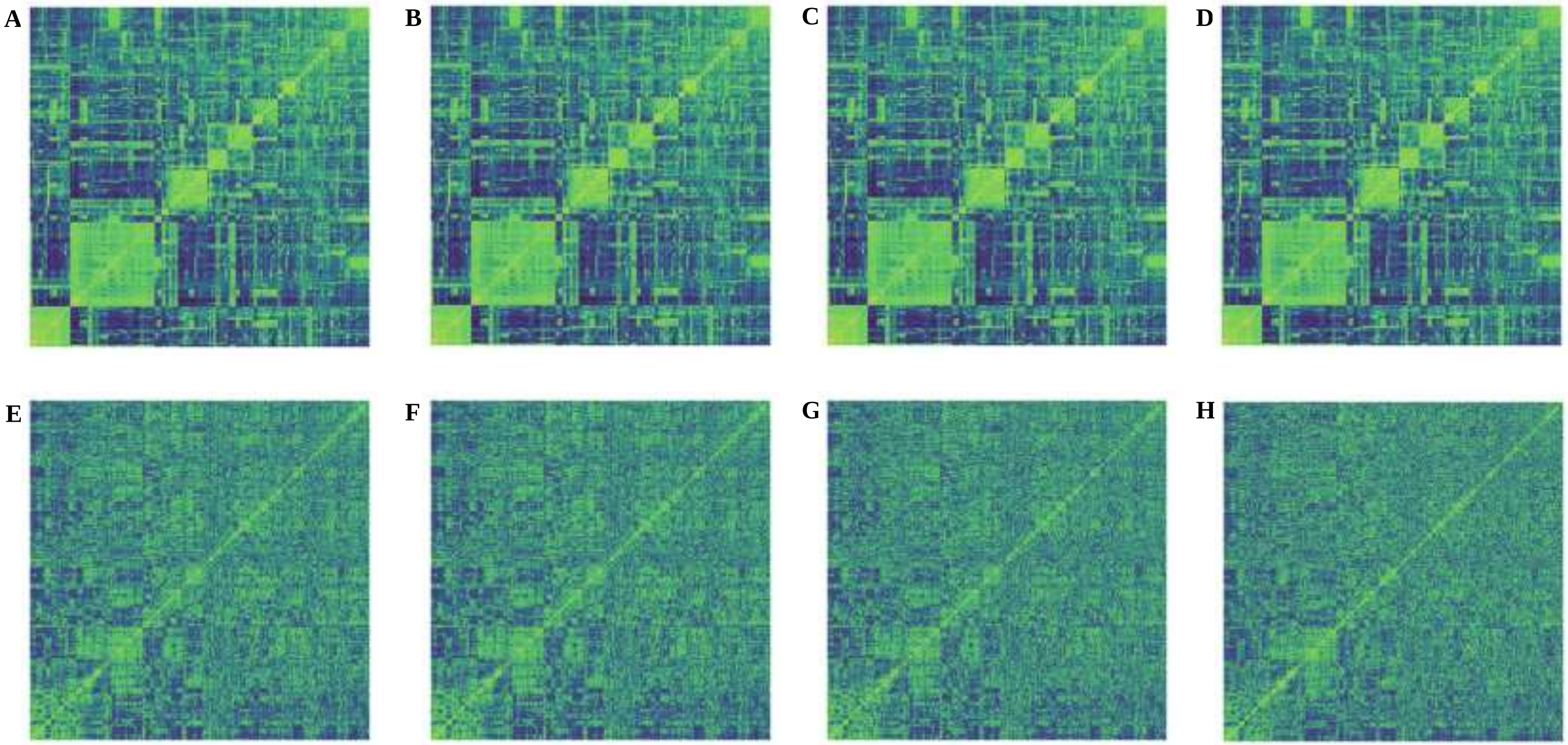


**Fig. S2 Some heatmaps for the additive (figures at the top) and dominance (figures at the bottom) kinships. A and E** Full-matrix (M_full) **B and F** reduced by 0.5 LD (M_0.5) **C and G** by 0.01 LD (M_0.01) and **D and H** optimized via algorithm for OTS (M_OTS), respectively. HEL dataset.
